# Identifying the Minimal Number of Protein Markers for Cell Type Annotation Using MiniMarS

**DOI:** 10.1101/2025.07.14.664825

**Authors:** Dhruti Parikh, Angli Xue, Hsiao-Chi Liao, Claire Wishart, Thomas M. Ashhurst, Givanna H. Putri, Fabio Luciani, Shalin H. Naik, Agus Salim, Felix Marsh-Wakefield, Raymond Hall Yip Louie

## Abstract

Over the past decade, there has been an explosion in the characterisation and discovery of cell populations using single-cell technologies. Single-cell multi-omics data, particularly those incorporating gene and protein expression, are increasingly commonplace and can lead to more refined characterisation of cell types. A common challenge for biologists is to isolate cells of interest using a minimal number of markers for cytometry experiments. Although several methods exist for marker selection, there is limited guidance on the relative performance of these methods, and a wrapper package that combines multiple methods is lacking. The method that performs best can vary depending on the dataset and it can be challenging for researchers to test multiple methods for a given dataset. To address these issues, we present MiniMarS (<u>Mini</u>mal <u>Mar</u>ker <u>S</u>election), an R package that serves as a wrapper for 10 different algorithms. It allows users to determine the best-performing algorithm for identifying the optimal number of markers that will delineate cell populations in their dataset. MiniMarS uses pre-annotated cells with protein features from CyTOF or sequencing-based assays such as CITE-seq and Abseq as input. Outputs include 1) the minimum number of protein markers required to identify the annotated cell populations using a range of marker selection algorithms, and 2) a range of metrics to evaluate the performance of each algorithm. MiniMarS effectively differentiated populations across various datasets, including those from human blood, bone marrow, thymus, mouse spleen, and lymph nodes, even after subsampling over 41,000 cells to 2,500 cells. MiniMarS also identified 15 markers from CITE-seq data, which were then used to successfully identify the same 11 cell subsets in a CyTOF dataset (F1 score>0.9). Additionally, we showed that by appropriately combining clusters, MiniMarS improves the F1 score of a rare population identification (<1% of total cells) by 28.7%. Together, these findings highlight the broad applicability of MiniMarS in identifying appropriate markers for distinguishing cell populations.

## Introduction

Single-cell technologies have enabled scientists to unmask the heterogeneity of complex biological systems at unprecedented resolution. In particular, single-cell RNA sequencing (scRNA-seq) captures transcriptional heterogeneity at the cellular level and has led to a rapid progression in the number of single-cell datasets available (Tang et al., 2009). For example, the technology has enabled the identification of rare immune cell populations and the discovery of new biomarkers (Nguyen et al., 2018; Van de Sande et al., 2023; Vandereyken et al., 2023).

Downstream analysis of scRNA-seq data involves grouping individual cells based on transcriptional similarity, enabling the identification of their cell type or state (Andrews et al., 2021). For downstream functional characterisation or isolation of these cell states, methodologies such as fluorescence-activated cell sorting (FACS) are commonly used, but these require cell surface protein expression. While protein expression can be inferred from gene expression, the reliability of these inferences is inconsistent, as gene and protein expression do not always positively correlate (Buccitelli & Selbach, 2020; Liu et al., 2016). Furthermore, many proteins are not expressed on the cell surface and cannot be used to isolate cell states using FACS.

The advent of single-cell multi-omics (Lee et al., 2020), such as technologies that measure gene and protein expression in the same individual (Peterson et al., 2017; Stoeckius et al., 2017), has further advanced our understanding of these cell states. It allows identification of cell types based on both gene and protein expressions. Simultaneous measurement of transcriptomic and proteomic information enables researchers to identify cell surface markers that discriminate transcriptionally distinct cells. Although 28 colours can be used on conventional cell sorters (Mair & Prlic, 2018) and up to 50 colours on spectral cell sorters (Konecny et al., 2024), these represent only a fraction of the markers that can be detected in single-cell experiments, which can range upwards of 200 proteins (Peterson et al., 2017; Stoeckius et al., 2017). Thus, a challenge for researchers is to identify the optimal set of protein markers that precisely distinguish cell subsets for isolation using downstream methods such as FACS.

Numerous computational methods (Pullin & McCarthy, 2024; Zappia et al., 2018; Zappia & Theis, 2021) have been developed to assist researchers in selecting markers that can distinguish cell types in single-cell datasets (Li et al., 2022; Missarova et al., 2021). However, each method can produce an inconsistent list of markers, with varying levels of performance determined using different metrics. Since the majority of methods have been developed for scRNA-seq datasets, researchers are left uncertain about the most appropriate methods and subsequently the cell surface protein markers to choose, hindering the reliable isolation of target cells.

To address these issues, we present MiniMarS (<u>Mini</u>mal <u>Mar</u>ker <u>S</u>election), an R package that serves as a wrapper for some commonly used single-cell marker identification methods, including CiteFuse (Wang et al., 2014), geneBasisR (Missarova et al., 2021), sc2marker (Li et al., 2022), non-marker specific algorithms and statistical tests such as XGBoost (Chen & Guestrin, 2016), F statistic, and five statistical tests implemented in Seurat (Hao et al., 2021) – Wilcoxon, bimodal, ROC (Receiver operating characteristic curve), t and LR (likelihood ratio) tests. The package utilises these established methods to identify the minimum number of cell surface protein markers required to distinguish one or multiple populations from each other. Furthermore, users can apply cross-validation to identify the best-performing method for their dataset using the implemented selection algorithm. The marker output from this algorithm can then be used to design antibody panels for FACS. MiniMarS provides visualisation functions to evaluate the performance of the chosen markers to differentiate the cell populations of interest. We perform extensive tests on single-cell datasets to demonstrate that our selection algorithm performs effectively. In summary, MiniMarS aims to use a selection-algorithm approach to aid scientists in identifying the minimal number of protein markers required to differentiate and isolate cell populations of interest in single-cell datasets.

## Results

### Overview of MiniMarS

The workflow of MiniMarS is illustrated in **Figure 1**. MiniMarS is an R package that identifies the protein markers that best differentiate pre-defined cell types or clusters. The first step of MiniMarS is to convert the normalised single-cell protein data matrix and corresponding cluster annotation to the desired format. The input can be: 1) a feature matrix with a vector of cluster labels, 2) a Seurat object, or 3) a SingleCellExperiment object. Clusters of interest are then selected, after which we sub-sample the data and divide it into training, validation and test sets (**Methods**).

**Figure 1.** MiniMarS Workflow. **A)** The first input is the feature matrix and associated cluster annotation, or a Seurat or SCE object containing this information. The second input is the clusters of interest that need to be delineated. **B)** Next, the data are subsampled for computing efficiency and then divided into training, validation, and test sets. **C)** The training set is used to identify the markers using 10 different methods. The validation set is used in a selection algorithm that combines the markers from the 10 different methods. The list of markers is then used with the test set to obtain the final performance measure, which is subsequently visualised.

The training set is first used for marker identification to determine markers that best differentiate the selected clusters (**Figure 1**). This is achieved initially by ten commonly used methods, including CiteFuse (Wang et al., 2014), geneBasisR (Missarova et al., 2021), sc2marker (Li et al., 2022), feature importance scores based on XGBoost (Chen & Guestrin, 2016), F statistic, and five statistical tests embedded in Seurat (Hao et al., 2021) – Wilcoxon, bimodal, ROC, t and LR tests.

The validation set is then used to measure the performance of the selected markers for each corresponding method (**Figure 1**). The performance measures are determined by first building a supervised machine learning model based on XGBoost, using the training dataset and corresponding cluster labels, where the matrix utilises only the predicted markers. Next, the validation dataset is passed through this model to predict the clusters. This prediction is then compared with pre-annotated clusters to calculate the precision, recall, and F1 score (**Methods**). Both unweighted and weighted scores are provided, the latter of which weighs the score based on the number of cells within each cluster. The performance measures are subsequently used for algorithm selection. There are two types of selection algorithms provided. The first selects the top N performing methods based on the F1 score, and the second selects all methods above a defined F1 performance threshold.

Finally, the test set is used for performance measurement of the ten methods and to select the best performing algorithm. The performance measures are calculated as described previously, but using the test set instead of the validation set. This procedure is repeated for a varying number (5, 10, 15, 20, 30, and 40) of markers, which are then used to determine the minimum number and identity of markers that best differentiate the defined clusters, subject to a minimum performance measure threshold. The performance for different numbers of markers can be visualised and validated using box plots. In addition, the overall results of the chosen markers can be visualised using UMAP, dot, and violin plots.

### MiniMarS is robust when using different numbers of cells or markers for training

As some of the marker-identification methods in MiniMarS require high computational power, the time it takes to run is dependent on the number of cells used for training. MiniMarS allows users to subsample the total dataset to reduce the runtime **(Supplementary Figure 1)**. To determine whether subsampling maintains high performance, we first applied MiniMarS to two human bone marrow datasets: 49,057 bone marrow-derived cells from six healthy controls (**Figure 2A, B, C**) and 31,586 cells from 15 leukemia patients (**Figure 2D, E, F)** (Triana et al., 2021). The healthy and leukemia samples were comprised of 13 and 14 broad cell types, respectively, based on 97 markers.

**Figure 2.**
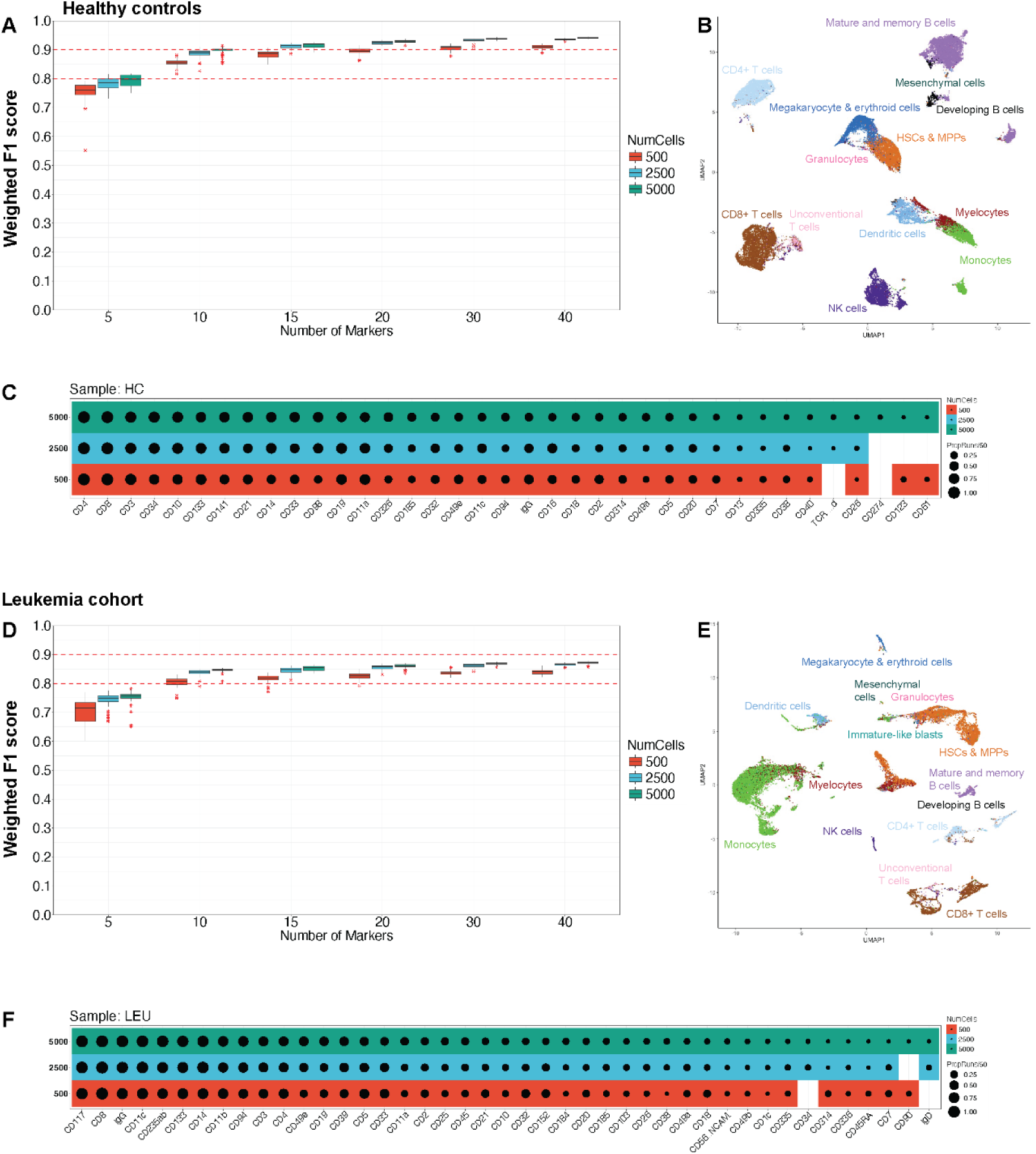
MiniMarS performs robustly across subsampled cells and varying numbers of markers. **A)** Boxplots of the weighted F1 score from the best-performing method across different numbers of markers, using 50 runs, with each run corresponding to a different subsampling of the total number of cells from the original dataset. Results shown for 500, 2500 and 5000 subsampled cells used for training. Horizontal red dashed lines show thresholds of 0.8 and 0.9. **B)** UMAP showing cluster locations, created using all cells and the 15 identified markers. **C)** The identified markers from each run, whereby dot size indicates the proportion of 50 runs where that marker was chosen. (A-C) are for the healthy dataset, (D-F) are for the leukemia dataset.

The original dataset was subsampled using 50 different random seeds. As expected, the weighted F1 score increased with the number of subsampled cells, reaching an F1 score of greater than 0.8 for 500 cells with 10 markers. We observed no substantial difference between 2,500 and 5,000 sub-sampled cells for both healthy (**Figure 2A**) and leukemia (**Figure 2D**) datasets. Most runs resulted in the same markers being chosen for 2,500 and 5,000 cells, for both healthy (**Figure 2C**) and leukemia datasets (**Figure 2D**), suggesting diminishing returns when subsampling under 2,500 cells.

When selecting 15 markers and sub-sampling 2,500 cells, MiniMarS provided a weighted F1 score greater than 0.9 for healthy controls (**Figure 2A**) and an F1 score above 0.8 for leukemia patients (**Figure 2D**). A UMAP plot using 15 markers for healthy controls (**Figure 2B**) and leukemia patients (**Figure 2E**) demonstrates that these markers are sufficient to distinguish the broad cell types. In contrast, using 5 or 10 markers leads to reduced differentiation between cell types (**Supplementary Figure 2**).

Based on these results, 15 markers yielded robust outcomes and were used as the default parameter in the MiniMarS package and subsequent analyses.

### MiniMarS improves the differentiation of rare cell populations using a cluster-combining strategy

A common application of cell sorting experiments is to ensure that rare populations are accurately isolated. However, markers for identifying rare cell subsets are implicitly weighted less in each method compared to markers that differentiate larger cell subsets. We thus investigated the performance of MiniMarS in distinguishing rare cell subsets and applied it to a combined dataset of cells from the mouse spleen and lymph nodes (Gayoso et al., 2021). This dataset comprised of 15,820 cells with 111 markers, from which we subsampled 5,000 cells. MiniMarS was then run using all subsets and 15 markers for differentiating eight broad immune cell types (**Figure 3A**). The unweighted F1 score for the two largest populations (B and T cells, > 90% of cells) was above 0.9, whilst the remaining six subsets (< 10% of cells) scored below 0.9 (**Figure 3B**).

**Figure 3.**
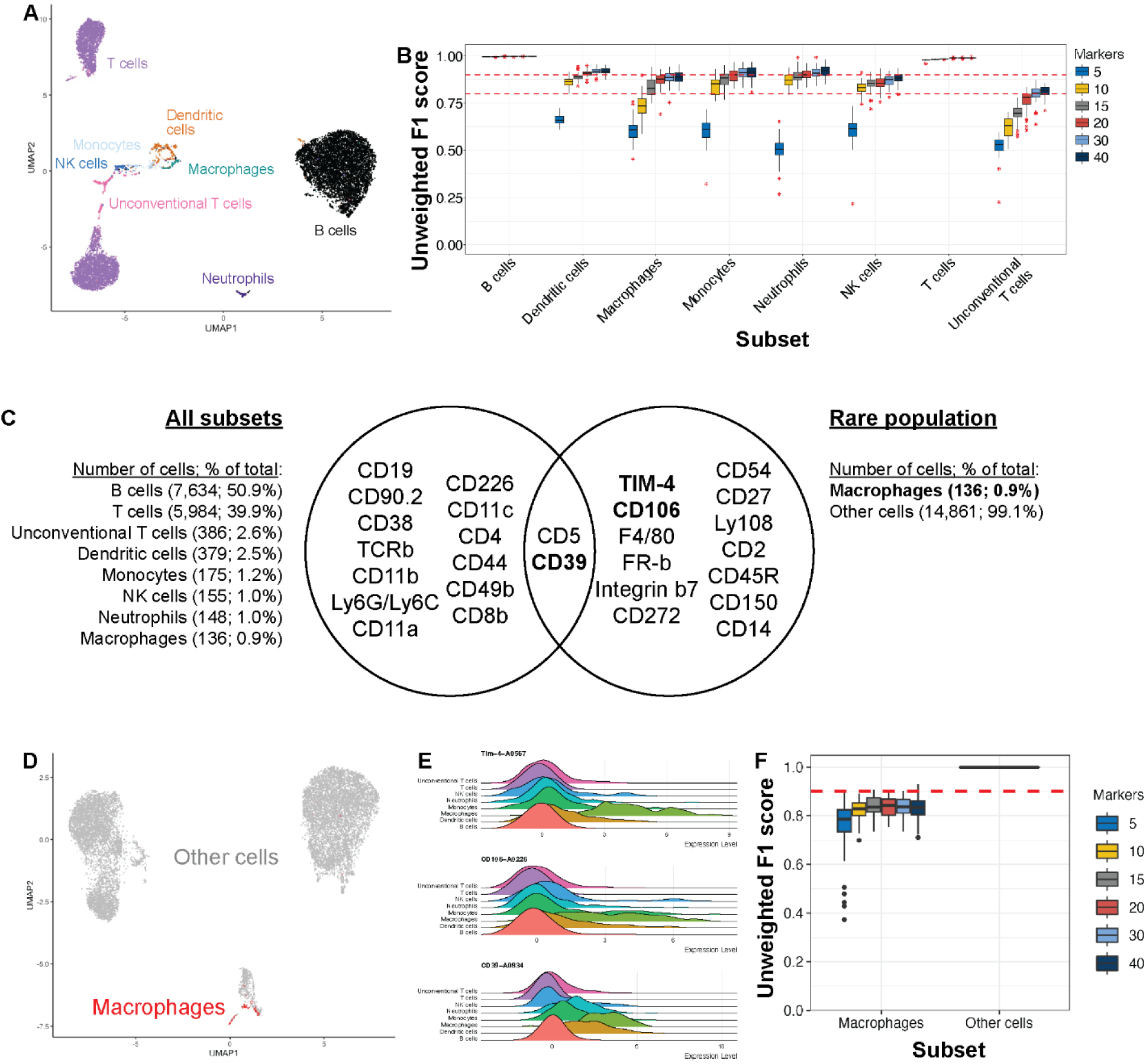
MiniMarS differentiates rare cell subsets. **A)** UMAP for the dataset using the top 15 selected markers, and **B)** The unweighted F1 score for each cell type using different numbers of markers. **C)** Top 15 markers for distinguishing all subsets (left) and the rare population (macrophages) from the remaining cell populations (right). The top two markers for macrophage differentiation are highlighted in bold. Numbers of cells (and percentages) are shown. **D)** UMAP generated by 15 rare population markers with macrophages in red. **E)** Marker expression for the top three markers for differentiating macrophages. **F)** Unweighted F1 score for macrophages when aggregating all other cell types as one, using different numbers of markers and sub-sampled to 5000 cells.

To test whether MiniMarS could improve the identification of rare cell subsets, MiniMarS was run by comparing the rarest subset (macrophages, < 1% of cells) to all other cell subsets combined. A Venn diagram illustrates the 15 markers selected when differentiating all subsets or focusing on the rare subset (**Figure 3C**). A UMAP using the 15 markers selected for the rare subset revealed a clear grouping of cells, with macrophages depicted in red (**Figure 3D**). Using ten markers, which include the known macrophage markers (TIM-4, CD106, CD39), it is possible to differentiate macrophages from all other cell subsets (**Figure 3E**) with an unweighted F1 score greater than 0.8. Increasing the number of markers to more than 10 has a minimal impact on performance (**Figure 3F**). There is a substantial performance gain when comparing the macrophage subset with the combined non-macrophage subsets (**Figure 3F**) rather than the individual non-macrophage subsets (**Figure 3B**). For example, there is a 28.7% increase in performance and an F1 score increase from less than 0.6 (**Figure 3B**) to approximately 0.8 (**Figure 3F**) when using only five markers. Together, these results highlight that MiniMarS has the capability to identify markers that separate rare subsets when an appropriate cluster-combining strategy is applied. The unweighted performance scores for different subsample sizes are shown in **Supplementary Figure 3**.

### Selected markers from MiniMarS can be applied across CITE-seq datasets

The results so far have used the same CITE-seq experiment for both training and testing. To examine whether MiniMarS performs well across samples from different experiments, two datasets from separate PBMC CITE-seq experiments were used as training and testing sets. The first “Hao” dataset comprised of 10,748 cells from eight patients, with 227 markers (Hao et al., 2021), whereas the second “COMBAT” dataset contained 3,703 cells from four patients, with 36 markers (Ahern et al., 2022). Two runs were performed, with one dataset used for training and the other for testing, and then vice versa. The performance of both runs achieved F1 scores greater than 0.9 (**Figure 4A**). 14 out of 15 markers were shared across datasets to delineate all cell populations (**Figure 4B**), indicating high reproducibility of the results, independent of the experiment. Finally, UMAP plots indicate a clear separation between clusters regardless of the dataset used for training or testing (**Figure 4C-D**). Dot plots show the relative marker expression for each subset (**Figure 4E**). The similarities between selected markers and performance across experiments provide confidence in the reproducibility of MiniMarS across experiments.

**Figure 4.** Validation of MiniMarS on training and test sets from two independent CITE-seq experiments. **A)** Weighted F1 score using the COMBAT and Hao trained datasets. **B)** Venn diagram showing markers in common between the two runs. UMAP plots showing **C)** Hao and **D)** COMBAT trained using 15 identified markers. **E)** Dot plots visualising marker expression across subsets.

### MiniMarS-selected markers are transferable across different experimental assays

We next provide a proof of concept that MiniMarS can identify markers using samples from a CITE-seq experiment, which can then be used to identify 11 cell populations in 160 samples from a CyTOF experiment involving 69,503 PBMCs (Ahern et al., 2022). The two datasets share 38 antibodies and 11 pre-annotated clusters. The UMAP plots generated from the 38 markers shared between both the CITE-seq (**Figure 5A**) and CyTOF (**Figure 5B**) datasets show a clear differentiation between cell populations. The key markers are verified by violin plots (**Figure 5C, Supplementary Figure 4**).

**Figure 5.**
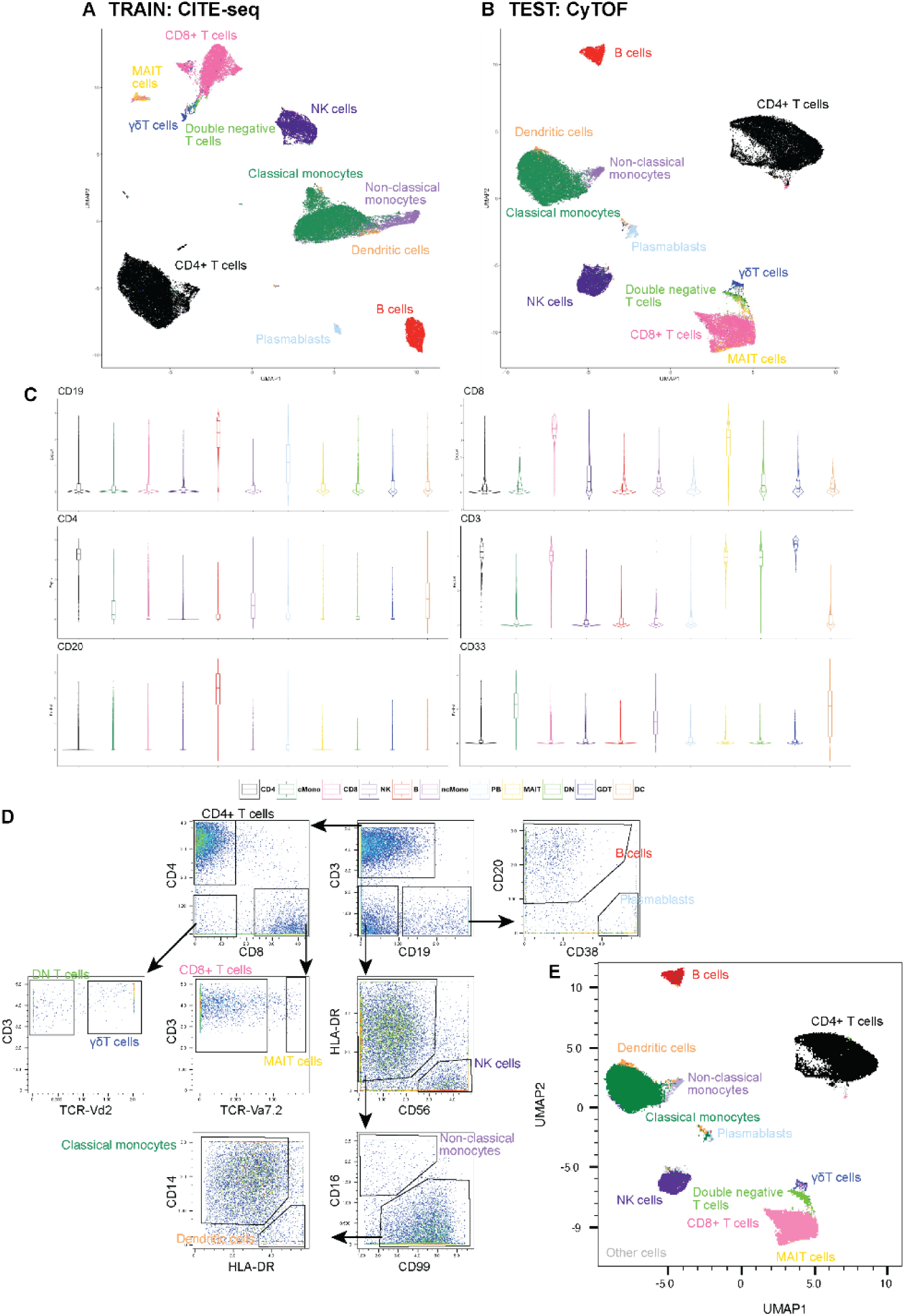
MiniMarS-selected markers are transferable on a matched CITE-seq/CyTOF dataset. **A)** UMAP of the training dataset using 15 markers identified by MiniMarS. **B)** UMAP of the test dataset using the same 15 markers. **C)** Violin plots from CyTOF using the top 15 markers (top 6 markers are shown). **D)** Manual gating strategy of the 11 subsets identified. E) UMAP using the manually gated cell populations (plus unidentified cells in grey).

The selected 15 markers from the CITE-seq assay were tested on the CyTOF dataset, resulting in a weighted F1 score of greater than 0.9 (**Supplementary Figure 5A**). This suggests that the markers selected by MiniMarS using the CITE-seq data are transferable. The 15 markers were then used to manually gate the 11 cell populations (**Figure 5D**), which are visualised in the UMAP plot (**Figure 5E**) using the CyTOF data, showing a clear separation of the cell types in the low-dimensional representation. The relative proportions of each subset were comparable across experiments (**Supplementary Figure 5B**).

### MiniMarS differentiates T cell states across a thymus dataset

MiniMarS was then tested on a more homogenous dataset, specifically T cells from a thymus dataset (Yayon et al., 2024). MiniMarS was run with 2,500 cells and 15 markers. It successfully differentiated T cell states (max weighted F1 score = 0.808) (**Figure 6A**). Identified markers are shown in **Figure 6B** and **Supplementary Figure 7**.

**Figure 6.**
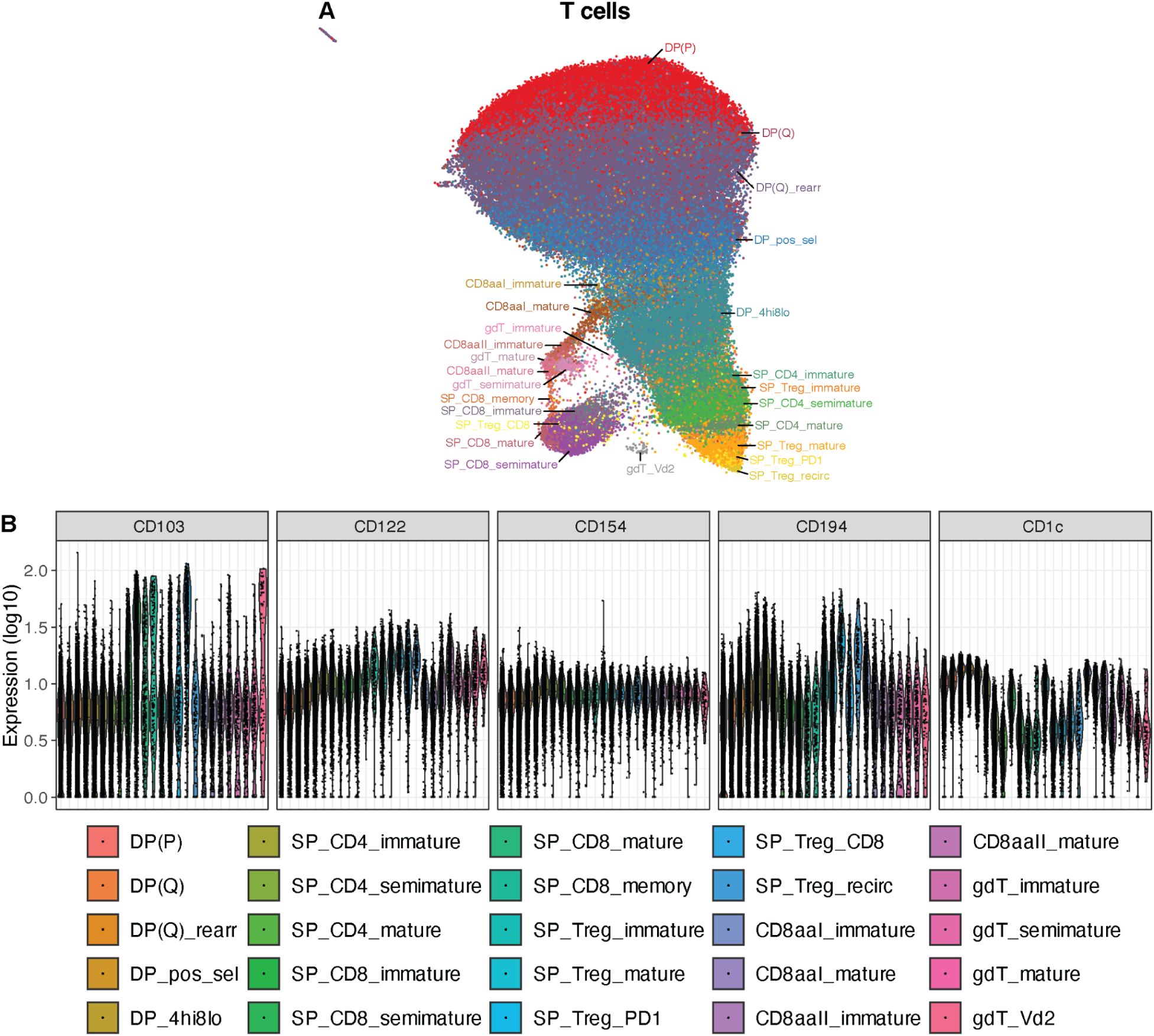
MiniMarS differentiates T cell states across a thymus dataset. MiniMarS was run on a thymus dataset. **A)** UMAP from MiniMarS run across selected CD4+, CD8+, T cells. **B)** Expression of five markers across T cell subsets.

## Discussion

Due to technological advancements, the number of protein markers that can be detected at the single-cell resolution continues to increase. Proteomics data is now commonly used with transcriptomic data to establish markers to distinguish cell states, which is crucial for downstream characterisation or isolation of cell subsets using cytometry experiments. While there are numerous methods available for marker gene or protein selection (Chen & Guestrin, 2016; Hao et al., 2021; Li et al., 2022; Missarova et al., 2021; Wang et al., 2014), their differing algorithms often result in inconsistent outputs, making it difficult to reliably identify cell-type defining markers to use for downstream experiments. To date, it is unclear how the performance of these different methods compares and the level of overlap in their output gene or protein marker lists. In this study, we introduce MiniMarS, a computational tool designed to identify the minimal number of protein markers for cytometry experiments by analysing single-cell gene and protein datasets. MiniMarS functions not only as a wrapper for existing single-cell marker identification algorithms, but it also implements method-selection algorithms to better integrate results.

Overall, MiniMarS offers a user-friendly interface that allows researchers to customise the marker identification process to suit specific experimental needs. We demonstrated the importance of utilising different methods to determine the optimal protein markers by implementing ten different methods in our wrapper. We provide sub-sampling options to reduce run-time and demonstrate that sub-sampling is robust and found that using 2500 cells and 15 markers for training provides satisfactory results. This feature allows the method to be applied to atlas-scale datasets. Furthermore, single-cell analysis has led to the discovery of rare cell subpopulations, and our wrapper has robustly identified markers for these rare populations. In particular, the top three markers that could differentiate macrophages from other populations were markers previously reported to be expressed by macrophages. These were TIM-4 (Wang et al., 2023), CD106 (Rice et al., 1991) and CD39 (Cohen et al., 2013). This shows the ability of MiniMarS to identify markers correctly, but the algorithm can also identify novel markers. In addition, MiniMarS successfully differentiated between different T-cell states within the thymus. We have also validated the effectiveness of MiniMarS by successfully identifying cell type-specific markers across various datasets, including a CITE-seq-CyTOF dataset, demonstrating its practical utility. For users with a pre-optimised sorting panel such as Optimised Multicolour Immunofluorescence Panels (OMIPs), it is also possible to calculate a performance score by inputting the selected markers.

Although we have validated MiniMarS across experiments and technologies, it must be noted that the optimal method or parameters depends on the dataset and the performance metric used. The performance metric can also influence which selection algorithm to choose. Thus, other performance metrics could be adopted in the future, and experimental validation of the selected markers should be undertaken. Future versions can incorporate additional marker selection methods to increase the robustness of the selected list of markers for researchers to use for experiments. Although MiniMarS might identify potentially novel markers, the algorithm’s consideration of a ground-truth set of markers is essential. Moreover, gene marker selection using MiniMarS can also be evaluated in future studies.

In summary, MiniMarS provides a crucial connection between the vast array of multi-omics single-cell experiments being generated. This link facilitates further research into exploring interesting cell states derived from these experiments.

## Methods

MiniMarS is a three-step pipeline designed for identifying and validating cell-type markers from single-cell protein data. The package is publicly available with detailed instructions found at (https://github.com/raymondlouie/MiniMarS). Below is a detailed breakdown of each step, including the methods used for marker selection.

### Step 1: Input Processing

MiniMarS begins by taking two primary inputs:

1. A single-cell protein matrix (e.g., CITE-Seq, Abseq, CyTOF), where rows represent cells and columns represent protein expression levels.
2. Corresponding cluster annotations that assign each cell to a specific cluster (e.g., cell type or condition).

To accommodate data from diverse sources, MiniMarS supports three input formats:

1. Feature Matrix: A standard matrix where each row is a cell and each column is a protein, along with a vector specifying cluster numbers.
2. Seurat Object: An R-based object from the Seurat toolkit for single-cell analysis, which includes metadata and clustering information (Hao et al., 2021).
3. SingleCellExperiment Object: An object from the SingleCellExperiment package, commonly used in R for managing single-cell data with cluster annotations (Amezquita et al., 2020).

After loading the data, MiniMarS selects a subset of clusters of interest for marker identification, allowing researchers to focus on specific populations. The data are then sub-sampled and split into training, validation and test sets. During this step, the user can choose between two strategies to subsample the training dataset: equal and proportional (default). ‘Equal’ subsampling will maintain the same number of cells for all clusters and favours rarer populations, whilst ‘proportional’ subsampling will maintain the same cluster proportions as the original dataset, resulting in an overall better performance across all clusters. For rare clusters, at least 4 cells will be retained.

### Step 2: Marker Identification

The training data are analysed to determine markers that best differentiate the defined clusters. MiniMarS applies ten distinct methods to identify these markers, each offering unique strengths for distinguishing clusters based on protein expression patterns:

1. CiteFuse uses a random forest model to quantify the importance of each marker using the mean decrease in Gini index (Wang et al., 2014).
2. geneBasisR dentifies important markers by iteratively adding markers with the largest difference between a “true” graph constructed using all markers and a “selection” graph formed using the current marker selection (Missarova et al., 2021).
3. sc2marker identifies markers using a maximum margin index measure that considers the distance of true and positive negative cells to the maximum margin (Li et al., 2022).
4. XGBoost is an ensemble machine learning algorithm that constructs boosted decision trees that separate clusters based on markers (Chen & Guestrin, 2016). Important markers are calculated first for each decision tree by how much the marker split point improves the performance measure, weighted by the number of observations the node is responsible for. The marker importances are then averaged across all decision trees.
5. fstat (F-statistic) is based on an analysis of variance (ANOVA) and identifies markers by measuring the variance in protein expression between clusters relative to within-cluster variance. A high F-statistic indicates a marker with a strong discriminatory power.
6. Five other tests are implemented using standard statistical differential tests with Seurat, namely, Wilcoxon Rank-Sum Test, Bimodal Test, ROC Analysis, t-test and Likelihood Ratio (LR) Test (Hao et al., 2021).

Selection-algorithms are then implemented to combine the markers identified by the different methods. We implement two types of algorithms. The first performs a majority-vote on the top N performing algorithms. The second performs a majority vote on all algorithms that achieve a minimum level of performance, with the F1 score used as a default performance metric. The default selection algorithm selects only the top performing method.

To maximise the customisation of the toolset for users, we also provide a function, *performanceOwnMarkers()*, which can calculate the cell type annotation accuracy based on a user’s input marker panel. This will depend on the user’s preference or availability of the potential markers (or antibodies for sorting) in the lab. The function will output the prediction accuracy for each cell cluster in the same format as the main function, so the results are comparable to that generated by predicted markers.

### Step 3: Visualisation

MiniMarS then provides visualisation tools, such as UMAP, dot and violin plots, to validate the chosen markers. In addition, box plots are available for visualising performance.

### Data processing

For **Figure 2**, we used a public CITE-seq dataset that contains healthy and leukemia samples of human bone marrow (Triana et al., 2021). To examine the ability of MiniMarS to identify broad cell types in different sample conditions, we analysed both healthy controls and leukemia datasets separately. Training and testing were done on each dataset separately, i.e., two training models were created, one for each dataset. In addition, we normalised the dataset using the CLR (Centered Log-Ratio) transformation, correcting for variations in protein sequencing depth across cells, and then scaled the proteins at the protein-level. The R functions Seurat::NormalizeData and Seurat::ScaleData were used. We checked the batch effect in the normalised datasets, which was not problematic across batches (**Supplementary Figure 6**).

For Figure 3, we used a public CITE-seq dataset that contains blood and lymph node samples (Gayoso et al., 2021). We normalised this dataset using the same pipeline as the one used for the dataset in **Figure 2**. There were moderate batch effects (**Supplementary Figure 6**), but they were not overly problematic as reflected by the relatively high F1 scores for this dataset (**Figure 2**). The “megakaryocyte & erythroid cells” subset contained 42 cells, which was too rare for the subsampling strategy, so this population was removed. As such, “macrophages” (consisting of 136 cells) were utilised as the rare population (**Figure 3**).

For **Figure 4**, we used two public healthy PBMC CITE-seq datasets, termed COMBAT (Ahern et al., 2022) and Hao (Hao et al., 2021). MiniMarS was trained on one random sample from the CITE-seq dataset and tested on the other, and vice-versa. The annotations provided in both studies were compared and used to generate a common cluster annotation, with 29 common markers identified. The protein counts provided by the authors were used as a direct input to MiniMarS, with no additional processing. Similarly for **Figure 5**, we used CITE-seq and CyTOF datasets from the COMBAT dataset (Ahern et al., 2022), with clusters for both sets re-annotated to ensure a common cluster annotation, with 36 common markers identified. The protein counts provided were used as a direct input to MiniMarS, with no additional processing. We trained one random sample of the CITE-seq dataset through MiniMarS, and the performance was tested on the CyTOF datasets. Manual gating was done on the CyTOF dataset using FlowJo v10.10 (BD Life Sciences).

## Acknowledgements

We would like to thank Hongjian Sun, Cindy Audiger, Rory Bowden, Mehrdad Pazhouhandeh, Jialel Gong, Yang Yang and Kelvin Tuong for their input to the manuscript and/or the initial design of the MiniMarS package. GHP and FMW are supported by the International Society for the Advancement of Cytometry (ISAC) Marylou Ingram Scholars Program.

