## Supplementary figures for "Identifying the Minimal Number of Protein Markers for Cell Type Annotation Using MiniMarS"

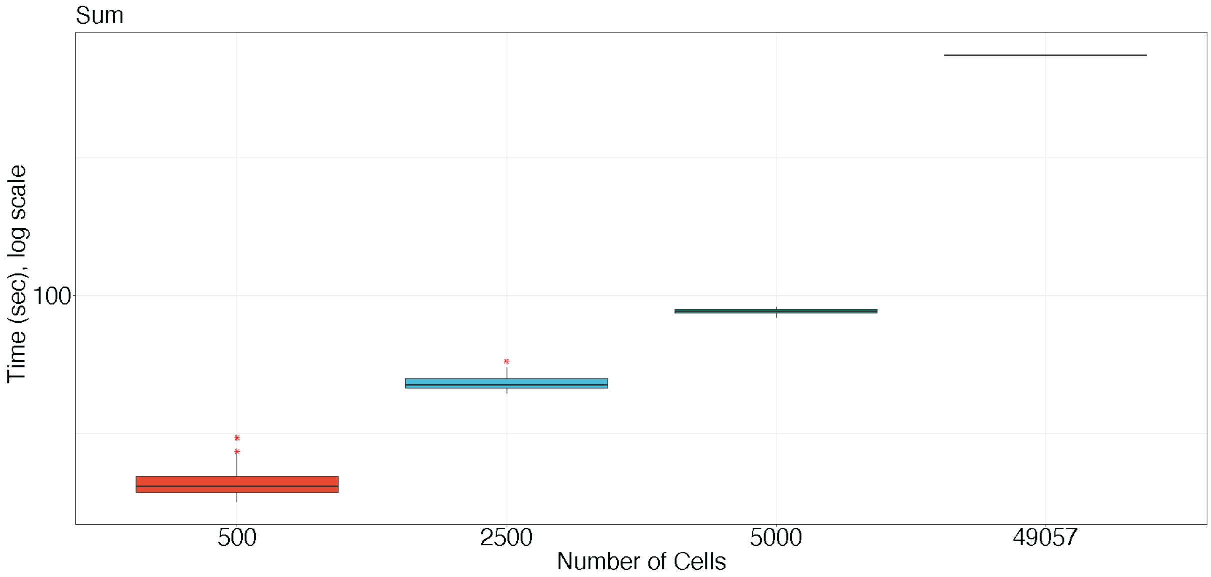


**Supplementary Figure 1.** Time to run all methods using different numbers of cells, including all 49,057 cells.

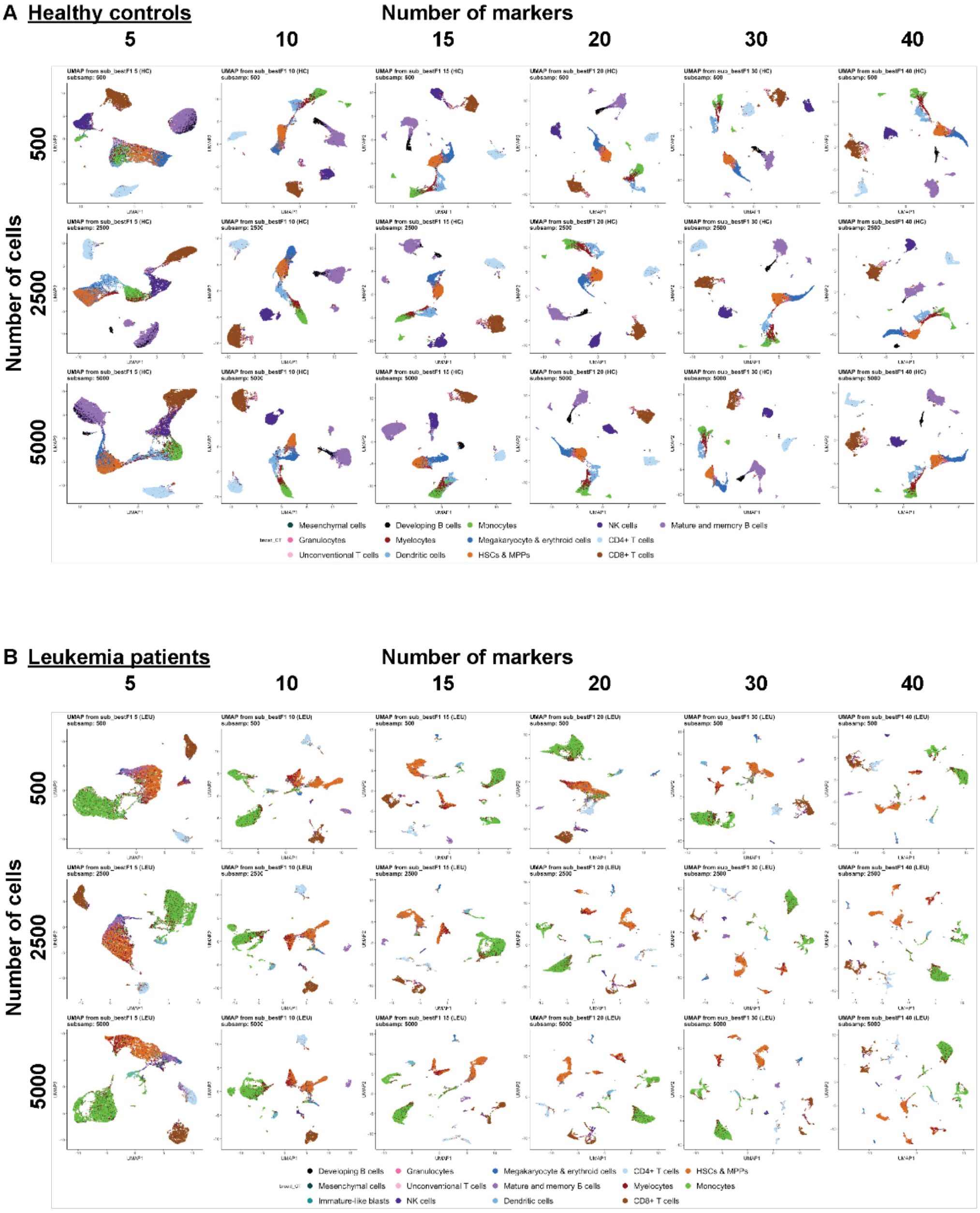


**Supplementary Figure 2.** UMAP plots of healthy **(A)** and leukemia **(B)** patients (from Figure 2) using different numbers of markers (columns) or cells (rows).

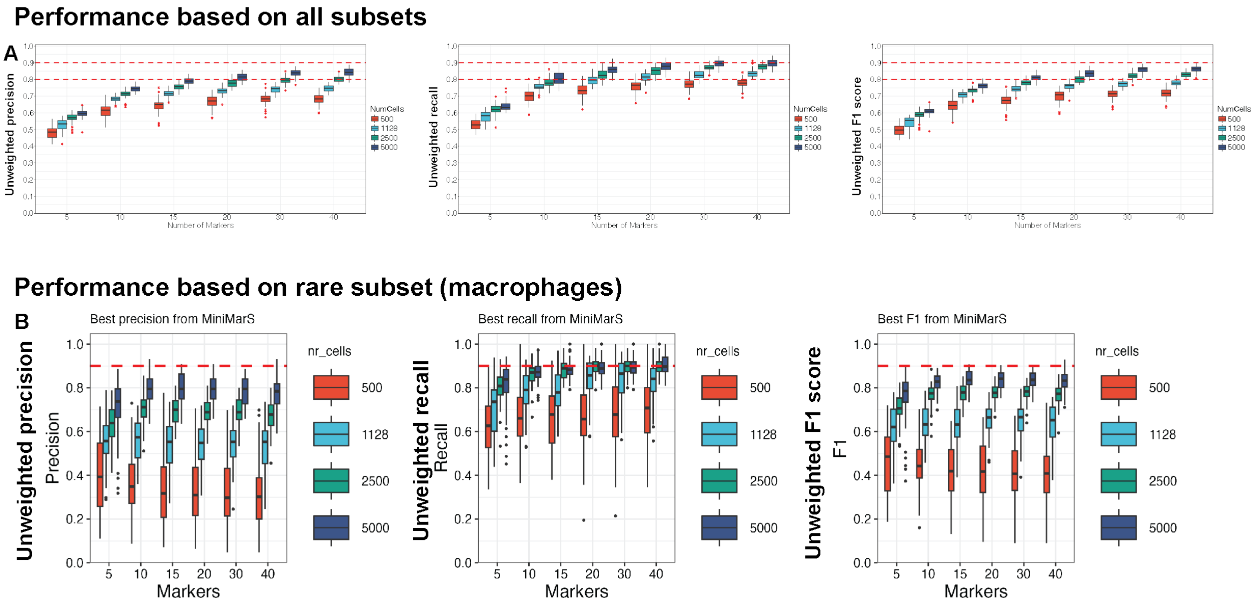


**Supplementary Figure 3.** Unweighted performance scores for differentiating between all cell subsets **(A)** or the rare macrophage population from all other cell types **(B).** Relates to **Figure 3**.

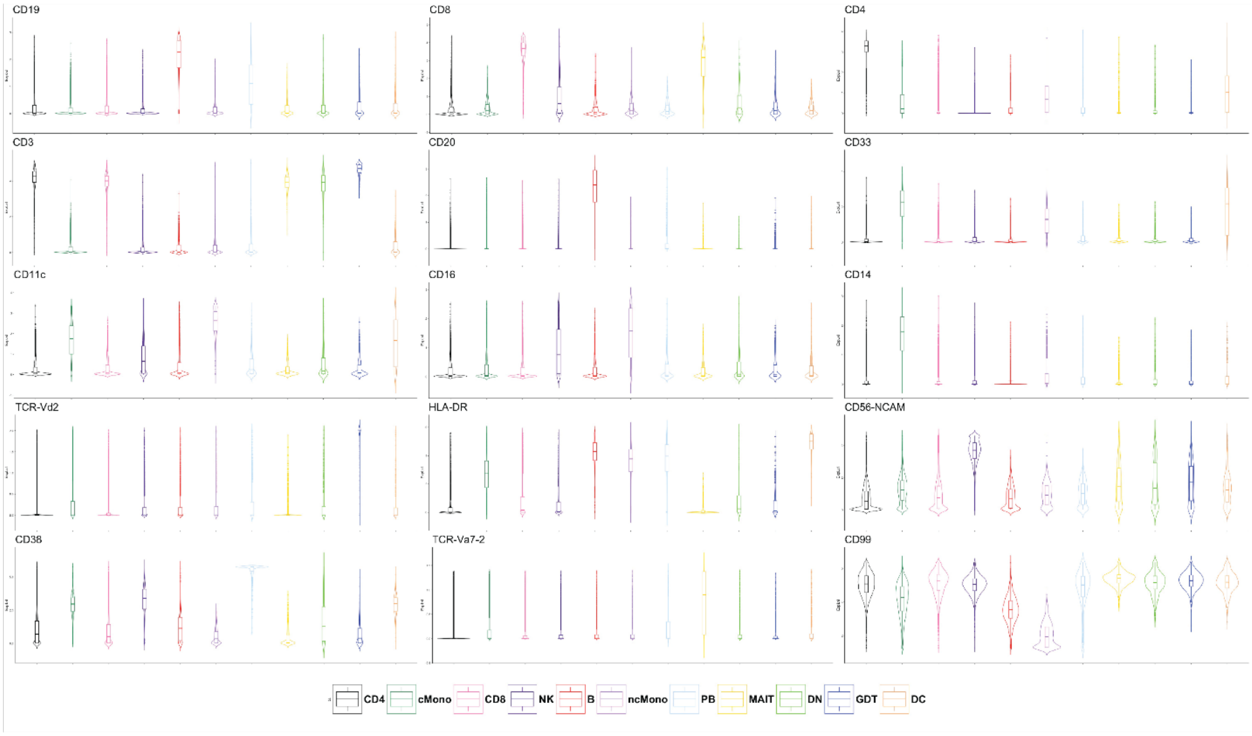


**Supplementary Figure 4.** Relative gene expression of oligonucleotide-conjugated antibodies across subsets in **Figure 5**.

**Supplementary Figure 5. (A)** Performance using CITE-seq dataset for training and CyTOF dataset for testing in Figure 5. **(B)** Proportions of subsets from CITE-seq (blue) and manually gated CyTOF (red) data.

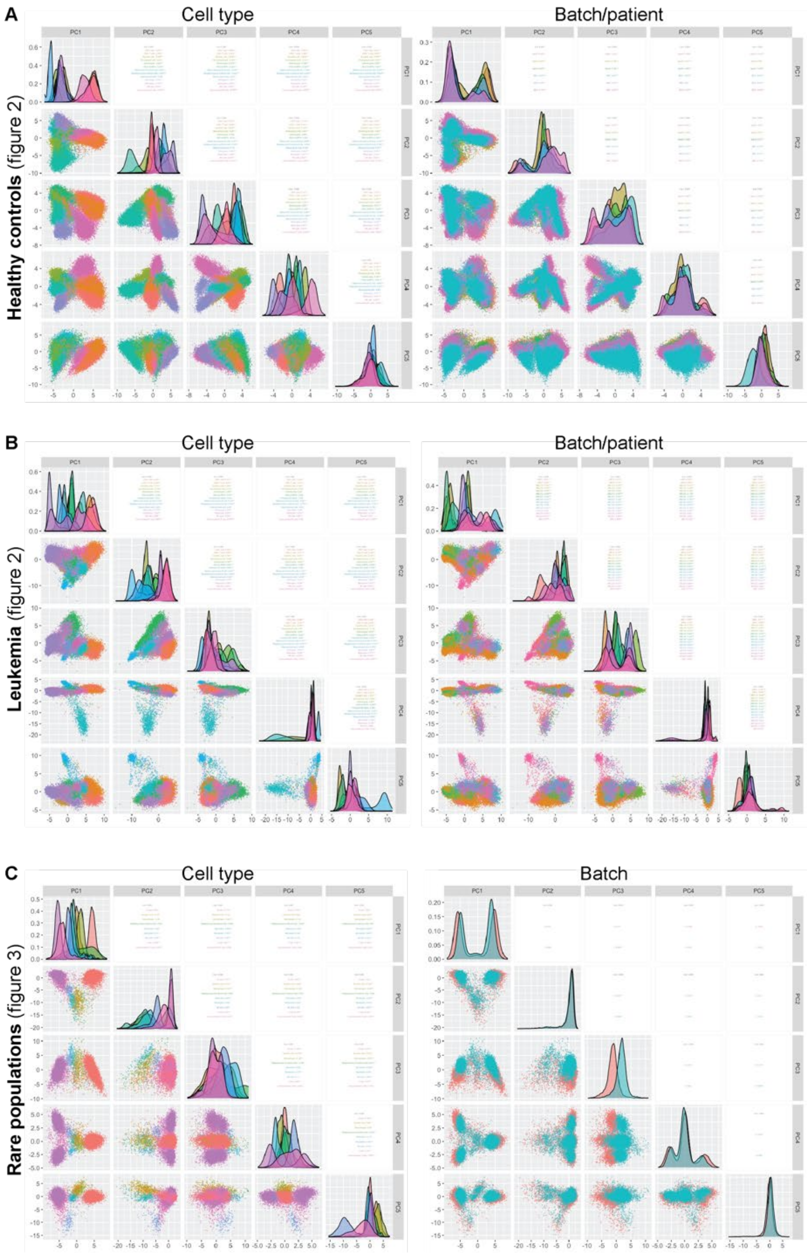


**Supplementary Figure 6.** Assessment of batch effects in **(A)** healthy controls, **(B)** leukemia samples, and **(C)** rare cell populations, with cells coloured by cell type (left) and by batch (right).

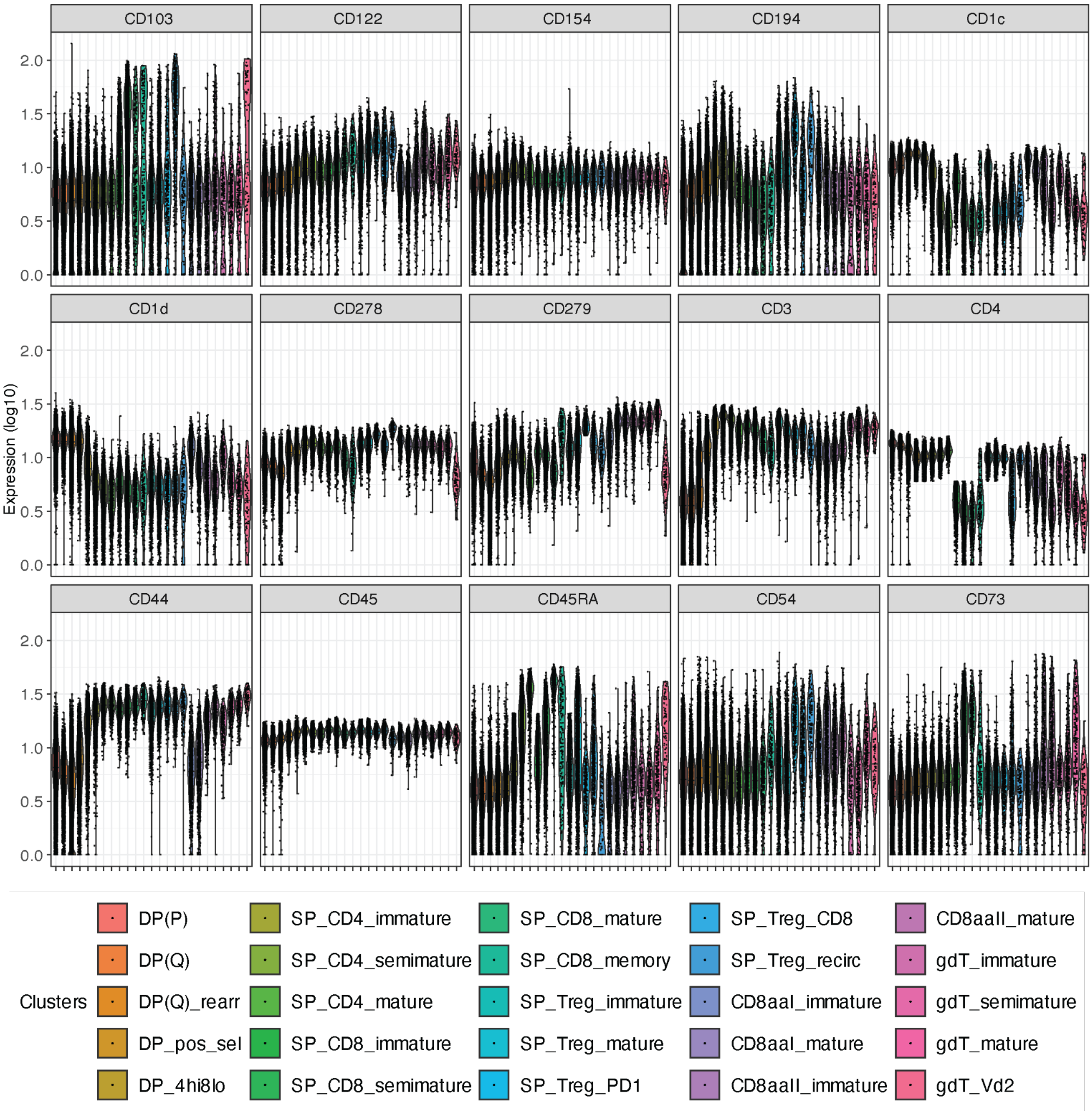


**Supplementary Figure 7.** Relative gene expression of oligonucleotide-conjugated antibodies across subsets in **Figure 6**.
